# Resurrection genomics provides molecular and phenotypic evidence of rapid adaptation to salinization in a keystone aquatic species

**DOI:** 10.1101/2022.07.22.501152

**Authors:** Matthew J. Wersebe, Lawrence J. Weider

**Affiliations:** Program in Ecology and Evolutionary Biology, Department of Biology, University of Oklahoma; 730 Van Vleet Oval, Richards Hall 304, Norman, OK 73019

**Keywords:** *Daphnia*, Salinization, Whole Genome Sequencing, LC50, Rapid Adaptation

## Abstract

Ecologists and evolutionary biologists are increasingly cognizant of rapid adaptation in wild populations. Rapid adaptation to anthropogenic environmental change is critical for maintaining biodiversity and ecosystems services into the future. Anthropogenic salinization of freshwater ecosystems is quickly emerging as a primary threat, which is well documented in the northern temperate ecoregion. Specifically, many northern temperate lakes have undergone extensive salinization because of urbanization and the associated increase in impervious surfaces causing runoff, and the extensive use of road deicing salts (e.g., NaCl). It remains unclear if increasing salinization will lead to extirpation of species from these systems. Using a “resurrection genomics” approach, we investigated whether the keystone aquatic herbivore, *Daphnia pulicaria*, has evolved increased salinity tolerance in a severely salinized lake located in Minnesota, USA. Whole genome resequencing of 54 *Daphnia* clones from the lake and hatched from resting eggs that represent a 25-year temporal contrast demonstrates that many regions of the genome containing genes related to osmoregulation are under selection in the study population. Tolerance assays of clones revealed that the most recent clones are more tolerant to salinity than older clones; this pattern is concomitant with the temporal pattern of stabilizing salinity in this lake. Together, our results demonstrate that keystone species such as *Daphnia* can rapidly adapt to increasing freshwater salinization. Further, our results indicate that rapid adaptation to salinity may allow lake *Daphnia* populations to persist in the face of anthropogenic salinization maintaining the food webs and ecosystem services they support despite global environmental change.

**Significance Statement:** Rapid adaptation to human-induced environmental change is critical for preserving biodiversity and ecosystem services into the future. A key question is whether populations of keystone species can rapidly adapt to maintain the ecosystems they support. We investigated rapid adaptation to anthropogenic salinization in *Daphnia pulicaria*, a keystone aquatic herbivore in lake ecosystems. By resuscitating decades-old resting eggs, we investigate genomic changes across an approximately 25-year temporal contrast from a severely salinized lake. We report that the genes showing signatures of natural selection throughout the genome are related to osmoregulation and ion regulation. Phenotyping clones for salinity tolerance revealed evidence that genetic changes may underlie rapid evolution. We provide molecular genomic and phenotypic evidence for rapid adaptation to salinity in *D. pulicaria*.

## Introduction

Ecologists and evolutionary biologists now recognize many examples of wild populations rapidly evolving in the face of environmental change (1, 2). A population’s ability to rapidly evolve is critical for survival in the face of ever-increasing anthropogenic environmental change. This capacity is especially important for organisms that provision key ecosystem services or are keystone species because their extirpation would fundamentally alter ecosystem dynamics. Despite this, studies demonstrating a mechanistic basis for rapid adaptation that integrates information from the genome to the phenome of a population are rare (3). A key reason for the paucity of such studies is that many loci of small effect are thought to contribute most to rapid evolutionary change and do not align with classical “hard-sweep” models making their identification difficult (4, 5). Additional analytical challenges are exacerbated by different population-specific parameters (e.g., effective size or N_e_) that influence the supply of new potentially beneficial mutations (θ = 4N_e_μ) and the resulting distribution of fitness effects (N_e_*s*) (6, 7).

One way that rapid adaptation can be studied is using temporal genomic contrasts. Most commonly, temporal contrasts take the form of so-called “evolve and re-sequence studies” which follow a population across time during experimental evolution trials that typically employ contrasting selection regimes (8–10). By finding the alleles that increase in frequency rapidly in different treatments, the molecular basis of phenotypic shifts can be explored (8). Such studies have been largely restricted to organisms such as bacteria (11) or *Drosophila* (10), which have rapid generation times and are easily manipulated in the lab. Other types of temporal contrasts encompass more natural experiments, such as the isolation and sequencing of ancient DNA, which can give insight into past selection (12). However, with few exceptions, the genomes sampled are divorced from the phenomes they produced; thus, inference is based solely on the change in allele frequencies. A third way that temporal contrasts can be studied is through “resurrection” studies that seek to hatch or germinate seeds, cysts or other resting stages of organisms and compare genotypes and phenotypes from different points in time (13–15). For instance, resurrection ecology (16–18), commonly used in animals from the freshwater crustacean genus *Daphnia* has provided insight into the genetic basis of various traits (15, 19, 20). Thus far, however, this method has not allowed the identification of loci that can be plausibly related to the phenotype under study because genetic markers are either too sparse (21) or the traits under study are too highly integrated across the genome (19).

*Daphnia* are keystone species in freshwater food webs, connecting the flow of energy from algal production to higher trophic levels such as fish (22, 23). Specific to many North American lakes, *Daphnia pulicaria*, maintains water clarity, a key ecosystem service and supports recreational fisheries with values in the millions of USD per lake per annum (24). Freshwater ecosystems are among the most threatened ecosystems worldwide, impacted by various anthropogenic stressors such as pollution, climate change and invasive species (25). One issue threatening many freshwater ecosystems is salinization due to human activities (26), within northern temperate lakes specifically salinization is particularly acute (27, 28). The causes and scope of salinization have been well known (29, 30), while more recent studies have focused on deciphering the ecological impacts of salinization (31, 32). In addition, we lack a more general understanding of this widespread environmental issue from an evolutionary perspective, and of the specific genetic architecture of adaptative responses. Such a perspective is critical because recent research has shown that current water quality guidelines do not sufficiently protect aquatic life from salinization (33).

To address this short coming, we sought to use resurrection ecology to study the evolutionary response of *D. pulicaria* from a severely salinized lake located in Minnesota, USA. Previous work on this lake has demonstrated the ecological dynamics of this population over the last 150 years (31). Towards this goal, we resurrected genotypes from across approximately 25 years from the sediment egg bank isolated from a dated sediment core. Using whole genome sequencing (WGS) of resurrected and extant individuals, we conducted numerous population genomic analyses to depict population structure over time, reconstruct the demographic history, and identify outlier genomic regions in the data. Additionally, we assayed a subset of genotypes for tolerance to salinity. Specifically, we were interested in testing two main hypotheses; first, we believed that the F_st_ outliers, or those genomic regions with extreme changes in frequency, would contain genes related to osmoregulation as selection would favor higher salinity tolerance. Second, this would be reflected in a higher mean tolerance of the more recent subpopulations. In addition, we identify a list of candidate genes, which likely influence phenotypic variation throughout the genome, and that can be targeted for further study.

## Results

We studied the *D. pulicaria* population of Tanners Lake (TL; 44°57’02.2”N 92°58’54.2”W), a small suburban hardwater lake located in Oakdale, Minnesota. The watershed includes approximately 32% impervious surfaces including parking lots, interstate highways and residential development (31). TL has received significant inputs of chloride from the watershed primarily in the form of the road deicer NaCl (524 kg Cl^-^ ha^-1^ yr^-1^) (34). The upper waters (i.e., surface/epilimnetic) chloride concentration of TL has increased significantly in the last 75 years, from approximately 1-2 mg Cl^-^ L^-1^ on average to over 150 mg Cl^-^ L^-1^(35, 36). TL has also transitioned to a state of cultural meromixis (37) with a persistent high salinity chemocline interrupting normal lake mixing dynamics. We isolated and sequenced 54 *D. pulicaria* clones including 10 from the water column and 44 resurrected from lake sediments representing an approximately 25-year temporal contrast (~1994-2019). In total we called 3,802,961 high confidence biallelic SNPs in the population.

### Population Structure and Genetic Divergence

Key to understanding the dynamics of the population across time is accurately describing population structure to rule out possible extinction and recolonization. The 54 clones selected for sequencing were separated into two clusters based on the first two principal component axes explaining 7.6% and 4.6% of the variance in the LD-pruned SNP data, respectively (Figure 1A). These two clusters largely separated the clones by depth with the older clones (layers 16-18 cm, 18-20 cm, and 22-24 cm) and more recent clones (Lake Clones, 2-4 cm, 6-8 cm, and 10-12 cm) forming the two groupings. Subsequently, we decided to assign the clones into three groups for the remainder of the analyses. We designated these as DEEP, encompassing all clones from 16-24 cm in depth (n = 18) and date from the mid to late 1990s. The second designation, called MID, encompassed all clones from 6-12 cm in depth (n = 18) and date from the mid to late 2000s. The third and final designation, referred to as TOP hereafter, included all clones collected from the water column and 2-4 cm in depth in the core (n = 18) spanning from 2016 to 2019. We based our assignment into the groupings on two observations: firstly, the TOP and DEEP subpopulations are delineated by the PCA clusters, and thus warranted separate assignments. Secondly, the MID subpopulation, while closely related to TOP, was intermediate in the time-scale –approximately 10 years prior to TOP and approximately 10 years after DEEP– and thus formed an appropriate intermediate grouping. While the TOP and MID are indistinguishable using PCA (Fig 1A), their temporal separation is a key feature of our study. Hence, we included each as a separate temporal deme in our simulations and analysis. Conveniently, this scheme also allowed each grouping to achieve equal sample size (i.e., 18 samples). Using Discriminate Analysis of Principal Components (DAPC), which is a flexible group assignment method, we found good analytical support for assignment of clones to the three *a priori* designated groups (Figure 1B). However, as with the PCA results, generally MID clones had non-negligible assignment probabilities when compared to the TOP subpopulation. Site-wise F_st_ was low across time, with an overall estimate of approximately 0.016 (Figure 1C). However, site-wise F_st_ estimates ranged from around 0 to as high as 0.398, with most sites having an F_st_ of essentially 0. The pairwise genetic distance between the three temporal subpopulations was related to the temporal distance with the TOP vs DEEP comparison being highest (0.019) and the MID population being intermediate to both (Figure 1D). Patterns of nucleotide diversity (π) were similar across all temporal subpopulations. Mean nucleotide diversity ranged from 0.0066 for DEEP, to 0.0070 for MID and 0.0069 for TOP (Figures S3-5).

**Figure 1:**
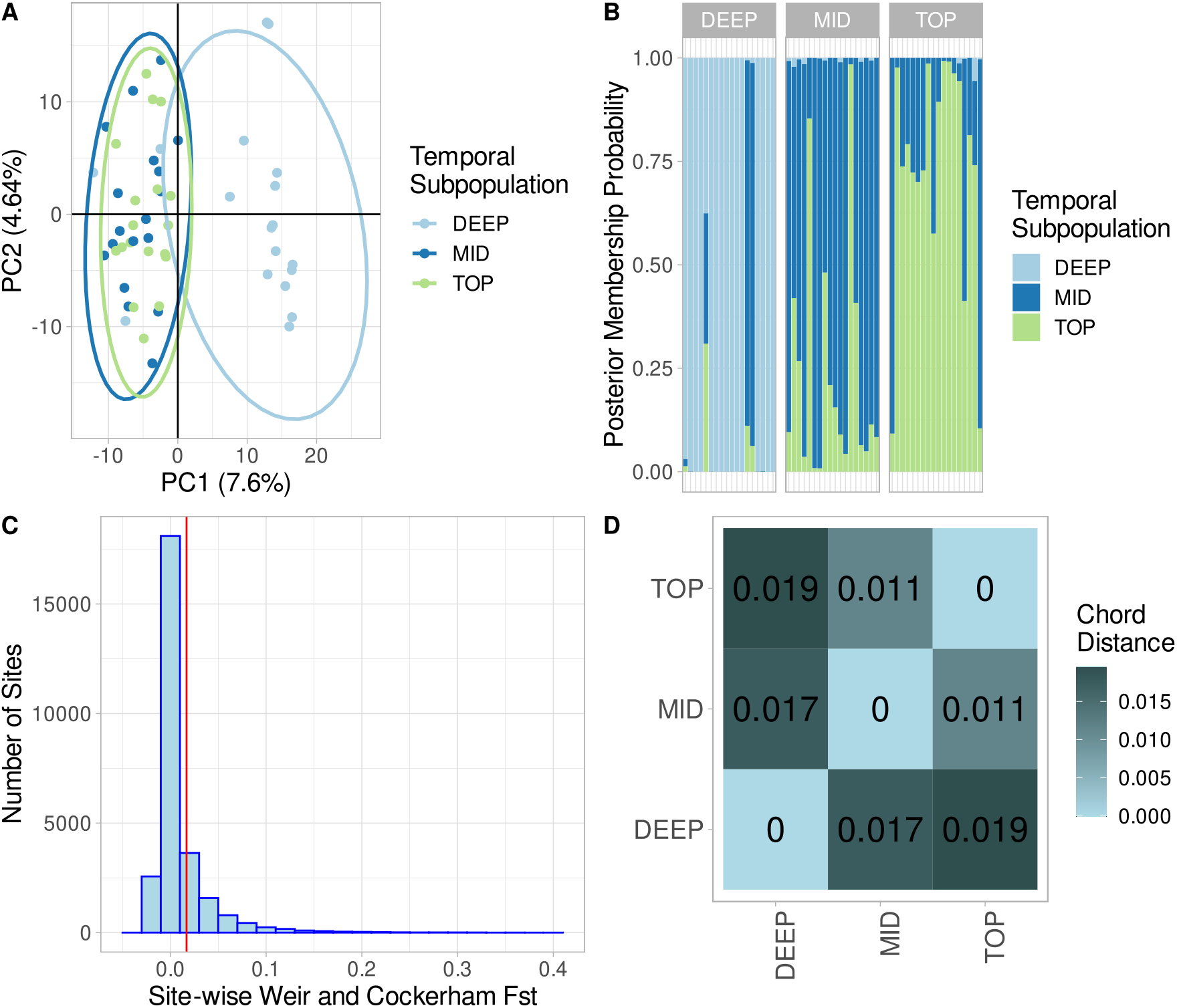
Genetic structure, divergence, and distance across time in Tanners Lake. A) Principal Component Analysis (PCA) biplot of all clones color-coded according to the depth of recovery in the core (cm). PC1 and PC2 explain 7.6 % and 4.6 % of variance observed in the SNP data, respectively. Older clones (22-24 cm, 18-20 cm, & 16-18 cm) form a cluster distinct from more recent clones (LC, 2-4 cm, 6-8 cm and 10-12 cm). B) Discriminant Analysis of Principal Components (DAPC) bar plot showing the posterior probability of assignment to the three *a priori* defined temporal subpopulations. C) Observed Weir and Cockerham site-wise F_st_ estimates. Overall F_st_ was low (red vertical line) at only 0.0169. D) Pair-wise genetic chord distances between the three temporal subpopulations (TOP, MID and DEEP)

### Estimation of Effective Population Size and Simulations

To accurately parameterize tests for selection, we sought to estimate a demographic model for the TL population. We were able to estimate the effective population size of the TL *D. pulicaria* population using both linkage disequilibrium (LD) and coalescent simulations. The LD-based results using just samples from the lake clone (LC) isolates showed that over the last few hundred generations, there was a period of population expansion and contraction (Figure S1). The population reached a peak N_e_ of 4000-6500 approximately 150 generations ago and the population has contracted recently to an N_e_ of around 2000. For this analysis, we interpret “generations” here to be sexual generations (i.e., LD is related to recombination and asexual generations are ameiotic). Since sex may occur once or at most a few times per year in stable lake habitats (38, 39), we interpret a single generation to be equivalent to one year. The coalescent-based FSC2.7 (40) run with the highest likelihood estimated N_e_ to be 2931 individuals and had a signal of population contraction with a population growth rate of 2.629 x 10^-5^. The estimate of effective population size was within the 95% confidence for 100 parametric bootstraps; however, the estimate for population growth rate was not and the confidence intervals included zero (table S1). We ran simulations in FSC2.7 using the maximum likelihood estimates to establish expectations for F_st_ based on the modeled demographic parameters. The results from 100 independent runs of FSC2.7 were pooled to develop a distribution of expected F_st_ values from simulated ~110000 SNPs. Testing each LD-pruned SNP against this distribution and correcting for multiple testing using false discovery rate (FDR) resulted in 178 outlier SNPs with corrected one-tailed p-values above a p = 0.05 significance threshold (Figure 2). There were outliers on every chromosome, ranging from a low of 2 SNPs (CHR 08) to a high of 42 (CHR 04).

**Figure 2:**
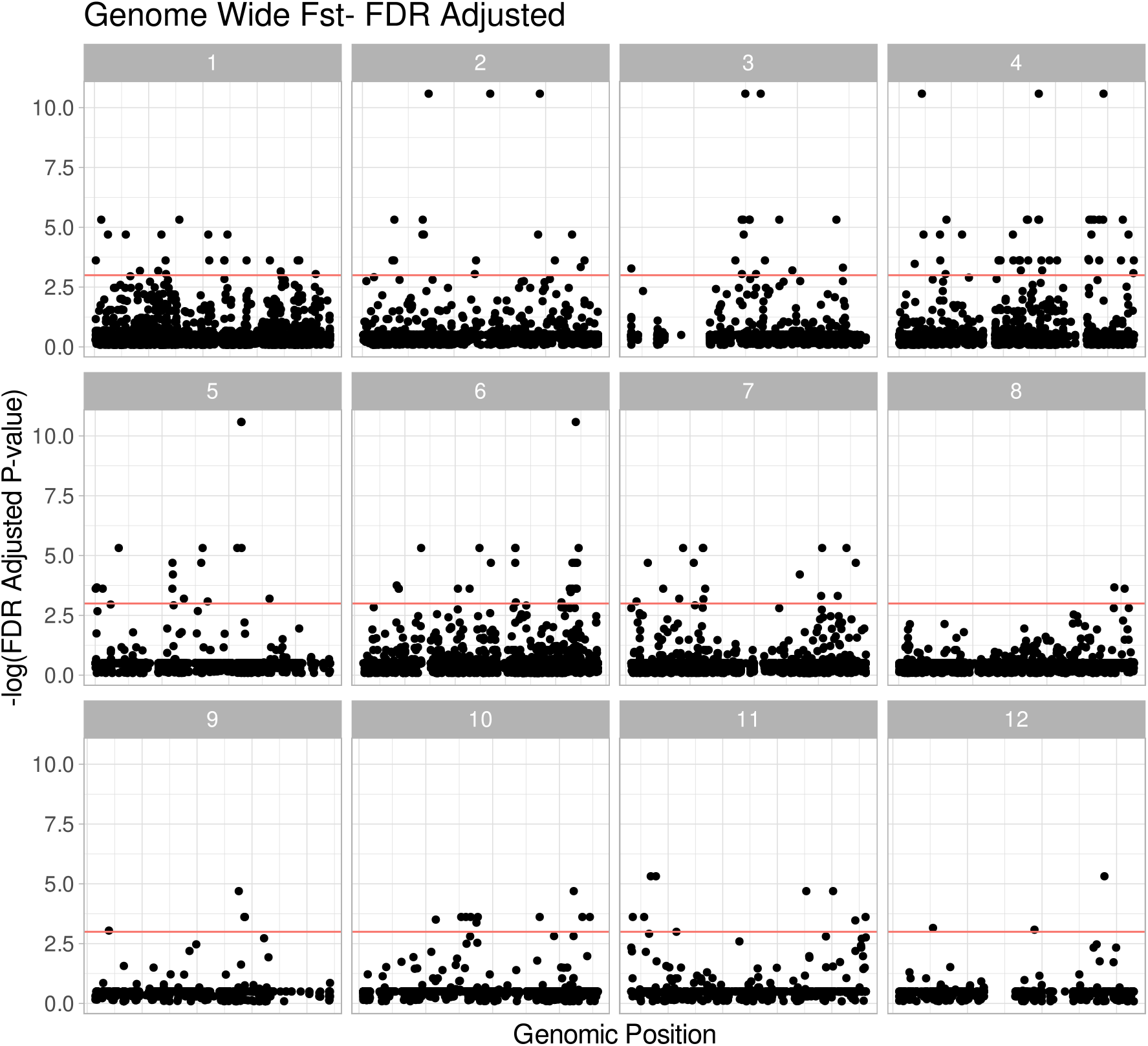
Genome-wide F_st_ Manhattan plots. Each panel represents a chromosome (1-12), each point designates a Single Nucleotide Polymorphism (SNP), the horizontal line indicates genome-wide significance at p = 0.05. Points above the red line are statistical outliers. Chromosome lengths are normalized across panels.

### Genes surrounding F_st_ outliers and GO term enrichment and Variant Annotation

One of our primary hypotheses was that the genes surrounding F_st_ outliers would be related to osmoregulation and salinity tolerance. We searched for genes within 10Kb of the F_st_ outlier SNPs (± 5 KB centered on the SNP) and in total we extracted 286 genes near these SNPs with known function in the *D. pulicaria* genome. GO term enrichment analysis with PantherDB webtool (41) yielded 59 enriched terms for this list of genes after correction for FDR (Figure 3). The enriched terms and p-values are available in table S2. Notable among the enriched terms for molecular function are chloride channel activity (GO:0005254; p = 0.00925); however, many different ion and channel terms were enriched. After running variant effect prediction (42) on all the SNPs called in the population, we identified 17181 variants with “high” predicted effects. Intersecting this list with outlier genes, we found 78 of the 286 outlier genes had high effect variants, only one of these, Chloride Channel 2 isoform X2 (*clcn2-x2*), located on chromosome 5 had any obvious relation to osmoregulation or salinity tolerance. *Clcn2-x2* has five SNPs of high effect, including four premature stop codon changes and a splice donor change that would likely severely interrupt protein function. It is tightly linked to the F_st_ outlier site at CHR05:12238, which is an intronic SNP within *clcn2-x2*. In addition to the high effect mutations, this gene has a total of 28 missense mutations and 37 synonymous mutations classified as moderate and low impact respectively. Of the clones surveyed for chloride tolerance (see next section), we found that the two most tolerant clones were homozygous for the wildtype (i.e., functional) allele at *clcn2-x2*. Overall, 252 of the outlier genes had effects that included moderate impacts to function such as a missense SNP.

**Figure 3:**
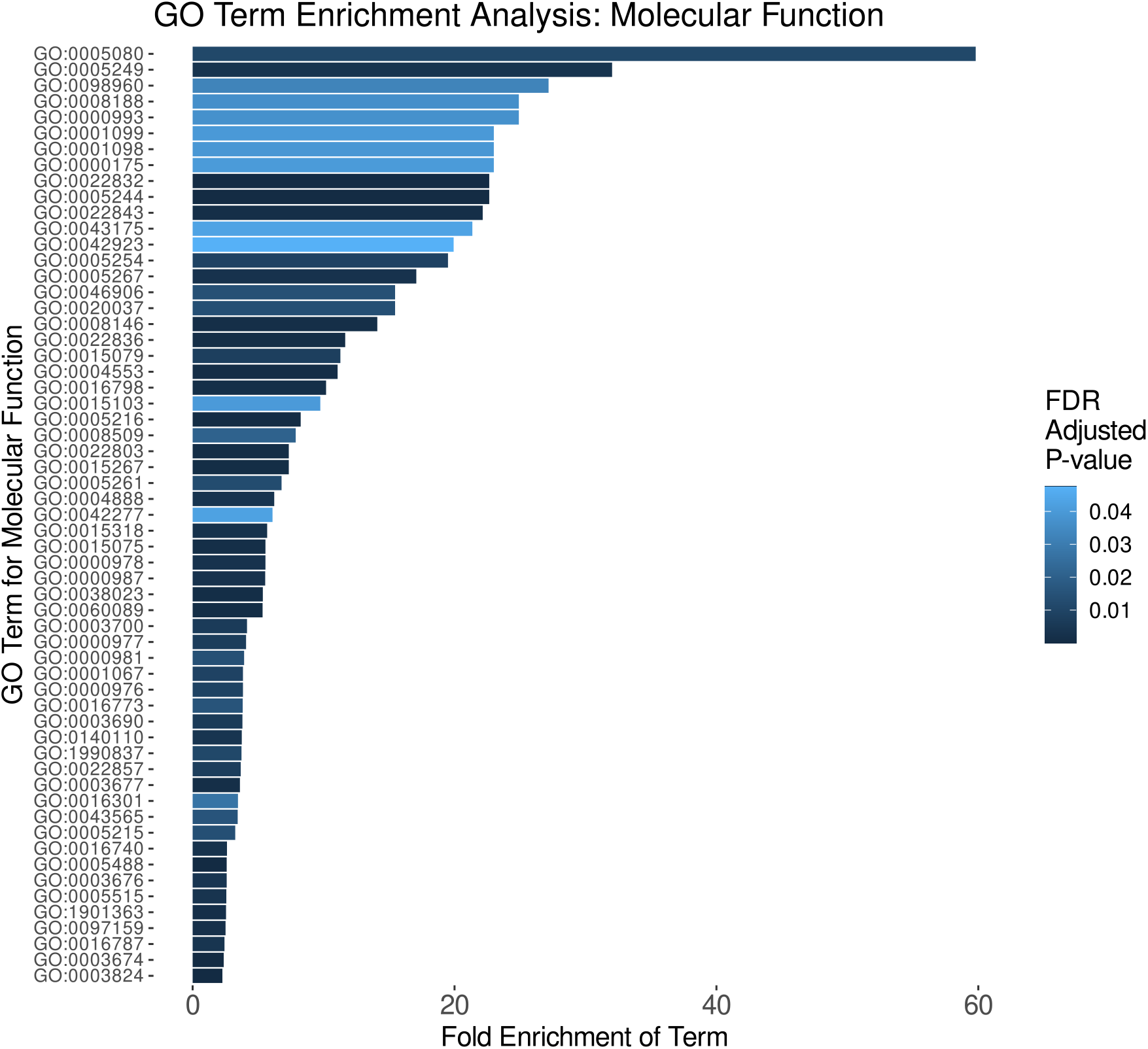
Enriched Gene Ontology (GO) terms for molecular function. Enriched terms are set along the Y-axis. The length of each bar indicates the term’s fold enrichment in the analyzed gene list, and the bar color denotes its False Discovery Rate (FDR) corrected p-value. In total, 59 terms were enriched among the 286 genes found near outlier SNPs. GO term mappings are provided in table S2.

However, *clcn2-x2*’s behavior as an outlier is not consistent with a single large effect locus. The SNP at CHR05:12238 is the 88^th^ most differentiated SNPs among the 178 outliers with an overall F_st_ of just 0.1985 (FDR corrected p = 0.0269). The top 11 outlier SNPs (F_st_ 0.3983 – 0.3175) all had F_st_ unobserved under neutral demography. One of the SNPs is an intergenic variant, while the 10 remaining SNPs are intronic SNPs associated with 13 different genes, 10 of which have annotated functions (table S3). The SNP located at CHR04:13394600 is the most differentiated SNP observed and is associated with the genes *pank4* and *tda6*. The remaining genes span across functional categories including the structural protein collagen alpha-1 chain and the calcium-binding protein *fstl5*, to several genes related to development comprising *still life* (43)*, daam1* (44), and *pnt* (45), which contains two outlier SNPs*, rap1gtp (46*), and *rotund (47)*.

### Chloride Tolerance LC_50_

We expected that increasing chloride pollution in TL would result in higher tolerance of more recent clones. Therefore, we conducted clone-specific assays to estimate 96-hour Lethal Concentration-50% (LC_50_). Clonal tolerance ranged from a low of 584.91 mg/L Cl^-^ (clones 10-12-12A & 18-20-04A) to a high of 1047.16 mg/L Cl^-^ for (Clone LC-06). We observed a main effect of subpopulation in the Kruskal-Wallis test (p = 0.006; Figure 4). Post-hoc testing using a pairwise Wilcoxon test found that the Lake subpopulation (i.e., TOP) was more tolerant on average than either the DEEP (22-24 cm & 18-20 cm) or MID (10-12 cm) subpopulations (p = 0.0076). The DEEP and MID subpopulations did not differ from one another (p = 0.9073).

**Figure 4:**
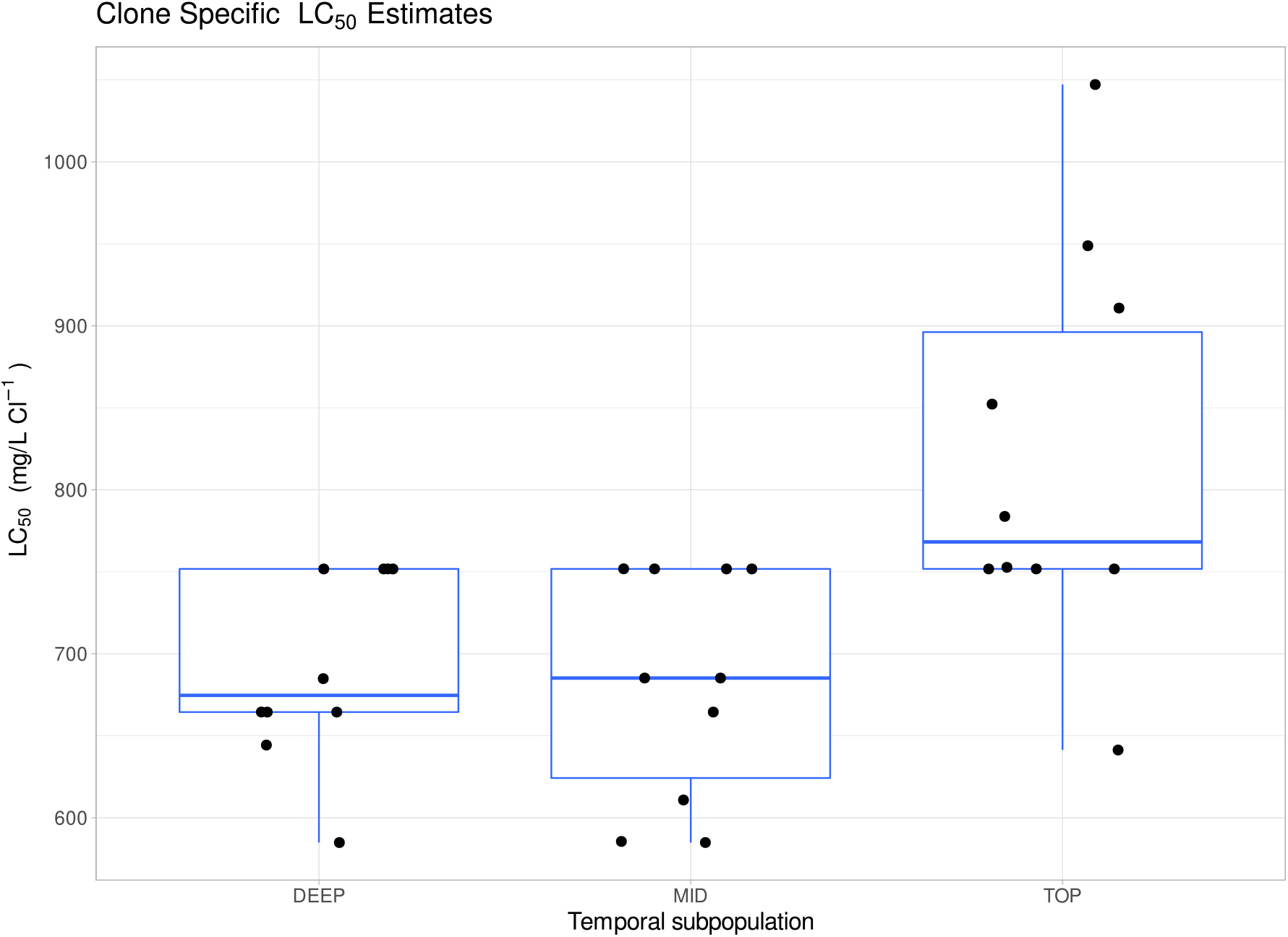
96-hour lethal concentration-50% assay (LC_50_) estimates for clones evaluated in the salinity tolerance experiments. Box plots indicate population medians and variances, each point is a clone. Overall, non-parametric ANOVA (Kruskal-Wallis) indicated a significant main effect of population. Post-hoc testing (pairwise Wilcoxon test) confirmed that the TOP population was more tolerant than either the MID or DEEP populations, which were not different from one another.

## Discussion

Our resurrection ecology (RE) study across ~25-year temporal contrast has provided phenotypic and molecular evidence of rapid evolution of salinity tolerance in the TL *Daphnia pulicaria* population. Using whole genome resequencing to analyze population genetic structure and demographic history, we have identified genomic regions putatively under selection, with support from LC_50_ chloride tolerance assays. We found that Chloride Channel-2 isoform X2 (*clcn2-x2*) has a unique mutational history and may be affecting salinity tolerance in this population, as one of many genes under selection in this population. We find support for both of our main hypotheses — genes related to osmoregulation are enriched in outlier regions and that the population shows increasing tolerance to salinity over time.

### Demographic History and Subpopulation Structure

A goal of our study was to establish a demographic model for the TL population. This would enable the modeling of population genetic summary statistics (i.e., F_st_) for establishment of a distribution of expected or null estimates of F_st_-based on the demographic history (discussed below). We used two complementary methods – LD-based methods and coalescent simulation to understand the recent demographic history of the TL population. Each method provides incomplete information, and each has unique biases that need to be considered. Furthermore, despite N_e_ being a critical parameter in population genetics, it is notoriously difficult to estimate (46). For instance, it appears that SFS (coalescent) based methods are underpowered for recent demographic history; however larger sample sizes may ameliorate some of these effects (47). This may also explain why it was difficult to accurately estimate the population growth rate parameter. The LD-based method we employed here appears to be strongly affected by the number of homozygotes in the sample (48), and thus, is sensitive to analyzing genomes from different generations. This prevented us from modeling each subpopulation separately. Additionally, it also appears to be inaccurate for the first few generations, giving unrealistically small numbers (see Figure S1). As such, we used these methods in tandem to increase the confidence in our estimation of the demographic parameters. Both methods were concordant that recent N_e_ was approximately 2000-3000 individuals, and both showed signatures of recent population contraction. This effective population size is not unexpected for *D. pulicaria* because populations are thought to delay sex and recombination for long periods of time (a year or more) (36, 37). During this time, clonal selection is thought to winnow down the population to a collection of ecologically equivalent clones (49). These facts, taken together with the current understanding of zooplankton metapopulation dynamics (50, 51) suggests that *Daphnia* effective populations should be small and insular.

Despite the small and insular nature of the TL population, it appears that drift does not always predominate. Our PCA results demonstrate at least on a decadal scale (i.e., between MID and TOP) that the population is temporally consistent. This likely reflects the dominance of the few ecologically equivalent clones predicted by Lynch and Spitze (48). It is likely that only after a rapid or pronounced change in environment can one expect to see a population structure across time, as reflected in other studies of *Daphnia* utilizing RE (15, 52). Salinization in TL is ongoing and likely started in the mid-20th century with the onset of widespread use of road deicing salts in the 1950s (53). The sediment intervals dating to the early 1990s from which we recovered the oldest samples in this study reflect a period of change in the sediment egg bank (e.g., low ephippial fluxes) (30) and suggests a period of rapid environmental change in this lake. Wersebe et al. (30), reported that by the early-1990s, TL had already reached a surface (waters) chloride concentration of at least 100 mg/L. Thus, all the source periods examined in the present study are typified by elevated chloride levels. Since we were unable to hatch eggs from before salinization commenced or was comparatively low in TL, it is impossible to determine if the structuring we observed was ongoing or more sudden. It is most likely that our temporal samples represent a process of ongoing adaptation to very rapidly increasing salinization. Regardless, since the clones are closely related across time at both the genomic and mitochondrial levels, it remains unlikely that the population became extinct and was recolonized by migrant genotypes (Fig. 1C; S2).

### Outliers and the Identification of Candidate Genes

The specification of a demographic model for the TL lake population allowed us to establish the presence of statistical significance for each site in the genome scan. This analysis revealed 178 regions with F_st_ beyond what we could reasonably expect based on demography alone. Genes surrounding these outliers had many functions, but as expected, genes involved in osmoregulation were among the most enriched in the dataset according to GO terms for molecular function. The presence of hundreds of outlier genes suggests that salinity tolerance has a complex genetic architecture and that tolerance to increased salinity requires changes at many loci of small effect. Transcriptomic studies of clonal isolates from within the *D. pulex* complex (to which *D. pulicaria* belongs) support this assertion (54). Indeed, cross-referencing these outlier genes with different mutational types (e.g., missense SNPs) showed that many of these genes have sites that may be under selection. However, the only gene with a known functional role in osmoregulation with high effect mutations was *clcn2-x2*. Chloride channels play a key role in osmoregulatory physiology of all animals. *Daphnia* are known as hypo-osmoregulators, meaning they attempt to maintain their hemolymph solute concentration above the ambient media concentration (55). Osmoregulation occurs within the gill-epithelium of *Daphnia*, and chloride channels like *clcn2-x2* play a major role in shuttling Cl- ions into the hemolymph across the basolateral membrane of the gill epithelium to maintain a hypo-osmotic concentration in the hemolymph (56). The constellation of premature stop codon mutations in this gene means that the protein is very unlikely to function properly because the channel is too short to pass through the cell membrane. Individuals are at least heterozygous for each of the four premature stop codon SNPs and the wild-type gene sequence annotated in the reference. Thus, they can produce a functioning protein. The two most tolerant clones in this study were both homozygous for the functional allele. This suggests that *clcn2-x2* is a critical gene requiring at least one functional copy and it explains some portion of the variance in acute salinity tolerance. However, a larger sample of genotyped and phenotyped individuals would be required to further validate this statistically.

The most differentiated genes in the genome were involved in a few key functions. There was a concentration of outlier SNPs in genes involved in regulating Rho-like GTPases which are involved in the regulation of actin filaments and neuronal development. The reason why these genes are the most differentiated is not initially clear. One potential explanation involves phenotypic plasticity. Increased salinity tolerance may require the accommodation of new developmental trajectories through phenotypic plasticity. Evolution via plasticity (59) requires that central developmental pathways serve as fuel for phenotypic differentiation (60). Furthermore, plasticity is thought to play an important role in rapid adaptation in *Daphnia*, a pattern we observe in the TL population (61–63). Regardless, testing of this assertion would be difficult because unlike other model arthropods, *Daphnia* development is not as well explored.

### Chloride Tolerance and Rapid Evolution

The clones observed in this study vary nearly two-fold in salinity tolerance (585 – 1047 mg/L), with the most tolerant clones detected in the most recent (TOP) temporal subpopulation. It is important to note that all clones in this study come from natal conditions that included elevated salinity with all subpopulations likely experiencing surface chloride conditions of 100-150 mg/L (31). Surface chloride conditions in TL are unlikely to exceed 150 mg/L in a year because of the dynamic equilibrium between annual loading and flushing of chloride in the system (49). However, TL more recently has transitioned to a state known as “cultural meromixis” were the accumulation of Cl at depth has interrupted normal lake mixing. For instance, in July 2019, we observed inferred Cl^-^ concentrations of approximately 275 mg/L directly above the chemocline (47% of the lowest LC50). Below the chemocline, inferred concentrations approached 483 mg/L (82% of the lowest LC50). Large-bodied *D. pulicaria*, undergo diel vertical migration to avoid visually oriented predators like fish. This vertical movement means that a clone may have to accommodate fluctuations in the ambient chloride concentration of 100s-of-miligrams over the course of 24-hrs. We attempted to study the degree to which this might occur in the summer of 2021 (see the supplemental materials for details). We conducted a Diel Vertical Migration (DVM) study in TL to track the spatial and temporal patterns of *D. pulicaria* distribution in the water column. We observed that *D. pulicaria* do inhabit the deepest, saltiest parts of the water column up to 12-m (Figure S6). However, in 2021, the chemical stratification of TL was much weaker than what we observed in 2019 and DO was not depleted at depth (see 31, Figure 1A-C & Figure S7 A-C). We believe the reason for the weaker chemical stratification is that the area surrounding TL (Washington and Ramesy Counties, MN) were classified as being abnormally dry on 06/29/2021 and had been classified as such intermittently since 09/29/2020. Thus, our abilities to determine the extent to which this might be reflected in a higher salinity year are diminished. Regardless, this shows that a portion of the *Daphnia* population is moving to more anoxic and saltier layers as part of the normal DVM behavior.

Such elevated Cl^-^ concentrations routinely observed in TL at depth are likely to have several sublethal effects that reduce clonal fitness. Presumably, a clone that has high acute tolerance has the physiological capacity to ameliorate the sublethal effects of increased chloride and we attempted to approximate this physiological response with LC_50_ assays. LC_50_ may be a quick measure of acute tolerance and it has an uncertain correlation with fitness components. As such, our phenotyping results should be interpreted with caution when considering the potential impact(s) on relative clonal fitness. Insofar as the pattern of rapid evolution of acute tolerance observed may be not actively reflect the phenotype actually under selection across time.

### Caveats, Analytical Roadmap, and Conclusions

All studies integrating historical data require caution to avoid over-interpretation of the emergent patterns. Our study is no different and relies heavily on a single core from a single lake. Our previous studies with this lake have explored many of these caveats (31). Specific to resurrection ecology (RE) studies, an acute limitation has always been hatching from the egg bank in the deepest layers. There may be non-random patterns of egg mortality in the sediments or non-random propensities in hatching success that skew phenotypic estimates and prevent accurate estimation of allele frequencies. Some have termed this the “invisible fraction” (*sensu* (50)). In our study this is most notable in our inability to hatch genotypes predating the most substantial increases in salinity (e.g., from 1950 or before). In other resurrection studies that have been able to hatch truly ‘ancient’ eggs (15), the utility of these samples in constructing a framework of phenotypic evolution is limited by sample size because only one or a few isolates survive to be cultured. In many circumstances, however, directly sequencing the eggs is an alternative but will prevent any possible phenotypic characterization of sequenced individuals because eggs are destructively sampled (51). These latter methods are still nonetheless difficult and involve costly genome amplification steps with variable success rates and high levels of exogenous contamination in the final libraries (51). This approach may help reduce the impacts of the ‘invisible fraction’ but will not eliminate it completely. Another caveat that we must highlight is that we have only analyzed a single population, and as such, we cannot place our results within a metapopulation context. We assume that gene flow should be negligible in producing allele frequency changes. To accurately account for gene flow, one would need to sample many additional spatially-distributed populations in addition to hatching temporal samples. While such data sets are within technical reach, the practicality of such a sampling regime would be difficult to amass for species such as *D. pulicaria*.

Regardless of the issues with RE data sets, we believe we can provide some insights into an analytical framework for future resurrection genomic studies. One major goal of the field of ecological and evolutionary genomics is to ultimately produce a comprehensive phenotype-to-genotype map-especially for those traits that one deems “ecologically relevant.” Relevant critiques of a QTL or QTN-centric research program aside (52), temporally-sampled genomic datasets may provide some very convincing examples of phenotype-genotype maps (see (53). Resurrection-type studies are poised to do this in natural populations as well. This will require robust demographic analysis of resurrected populations, integrated with simulations in a flexible manner. One potential way forward is the use of flexible-forward genetic simulations (e.g., those available in SLiM; (54)), which can allow more analytical power in genomic analysis of temporally sampled populations.

In summary, resurrection ecology (RE), when paired with whole genome sequencing, provides a unique and powerful way to study rapid evolution of populations *in situ*. Integration of candidate loci identified with RE into study designs including breeding experiments (55), forward mutation screens (56) or CRISPR technology (57) will provide unrivaled insight for the genetic architecture of complex, ecologically relevant traits in the wild. Such an approach would be beneficial here, for example in validating the effect of the null *clcn2-x2* allele on salinity tolerance in the TL population. Overall, however, here RE provides both molecular and phenotypic evidence of rapid adaptation in the TL population. Thus, the persistence of *D. pulicaria* in this severely salinized lake is likely the result of this rapid adaptation. Our findings indicate that keystone aquatic species such as lake *Daphnia* may continue to thrive in lakes that exceed current water quality limits for salinization and thus maintain the stability of the food webs and ecosystem services they support.

## Materials and Methods

### Clone bank and Sequencing

On 2 July 2019, we collected duplicate sediment cores in TL from a 14 m deep station following Wright (58). During the same period, we also collected *D. pulicaria* from the active plankton community using several vertical tows of a Wisconsin net at the core sampling station. Animals were isolated as single individuals in 125 mL plastic (screw-capped) cups in COMBO media (59). A total of 10 of these clones were established in laboratory culture. Resting eggs (ephippia) collected from throughout the cores were collected according to the methods outlined in Wersebe et al. (31). *D. pulicaria* are cyclically parthenogenetic, meaning they produce clonal offspring during the growing season and may occasionally engage in sex to produce resting eggs encased in durable structures called ephippia (60). Ephippia identified as *D. pulicaria* were subjected to hatching protocol described (15) (see supplemental). These hatchling individuals were expanded in culture to establish upwards of 10 clones per sediment layer to establish a clone bank. Fifty-four isoclonal lineages from the clone bank were selected for DNA extraction and whole genome sequencing (average 10X) on an Illumina NovaSeq by the Oklahoma Medical Research Foundation.

### Bioinformatics

Raw sequencing reads were quality trimmed and adaptor contamination removed using Trimmomatic (61). Quality trimmed reads were aligned to the chromosome-level *D. pulicaria* genome assembly (62) using the BWA mem algorithm (63). The resulting files were piped through samtools (64) to mark duplicates, fix mates and sort the bam files. We called variants using the bcftools mpileup and call pipeline using all individuals together (65). Using bcftools, the resulting BCF files were concatenated together into a single genome-wide file and quality filtered to a set of high confidence biallelic single nucleotide polymorphisms (SNPs).

### Population Structure and Genetic Divergence

The variants in the final quality-filtered VCF file were pruned for linkage disequilibrium (LD) using Plink (66) independent pairwise function (settings: 50, 10, 0.1), providing an independent and essentially random set of SNPs. The resulting variants were further filtered to SNPs with 0% missingness to a set 27854 genome-wide SNPs. We conducted Principal Components Analysis (PCA) in R (Version 4.2.0; R core team 2022) using the packages *adegenet* and *vcfr* (67, 68). We also conducted Discriminant Analysis of Principal Components (DAPC); a flexible population assignment method also implemented in *adegenet* (69). Using the cross-validation procedure outlined in the vignette, we found that retention of 10 PCs performed best in population assignment. We performed population assignment tests for each clone retaining 10 PCs and 2 discriminant functions and plotted the results as a bar plot to visual the probabilities of population assignment. DAPC is flexible enough to handle mixed clonal and temporal sampling - two factors that violate other assignment techniques (e.g., STRUCTURE (70). From the PCA and the DAPC, results, we determined that the samples could be assigned to three “sub-populations” according to the depth of their recovery (see results). Using these three subpopulations as designations, we estimated overall site-wise F_st_ using the basic.stats function and mean pairwise genetic distance genet.dist function in the R package *hierfstat* (71). In addition, we estimated nucleotide diversity (π) in 10 kb windows throughout the genome for each of the temporal subpopulations using the program PIXY (72).

### Estimation of Effective Population Size and Simulations

To parameterize our tests for selection, we sought to identify the recent demographic history of the TL *D. pulicaria* population. To accomplish this, we used two different methods to estimate effective population size (Ne) and growth trajectory of the population. The first method, implemented in GONE (73) uses LD to estimate the recent population history. This method is robust to non-equilibrium histories such as selection (74); however, it is suitable only for sample pools collected from the same generation. Thus, for this method we used only the samples collected from the water column in 2019. We ran six independent runs of this method using random subsets of 600,000 SNPs from all the SNPs called in the population and a constant recombination rate of 7.2 cm/mb estimated from the *D. pulicaria* genetic map (62). This analysis does not assume a given model *a priori*, instead it produces a population size trajectory that when inspected graphically can hint at different events (e.g., bottlenecks). In addition to LD-based methods, we estimated demographic parameters of the TL population - estimating N_e_ and the population growth rate - by fitting a demographic model to the folded site frequency spectrum (SFS) implemented in FastSimcoal 2.7 (FSC2.7) (40). With the LD pruned SNPs used above in the PCA analysis, we created folded 2-D SFS from the three temporal subpopulations. Using 100 independent runs of FSC2.7, we chose the run that maximized the likelihood of the observed data. We fixed the sampling points in time for the temporal subpopulation (MID and DEEP) to be 50 and 100 generations in the past. This assumes approximately 4 asexual generations a year and a single sexual generation for a total of five generations a year. Each run conducted 1-million coalescent simulations and used 40 brent maximization cycles. Further, we estimated empirical p-values for the site-wise F_st_ estimates using simulation (75). We chose the best fitting demographic model estimated in FSC2.7 and conducted 100 separate runs of this model to simulate approximately 1100 SNPs for each run. For each run, we estimated F_st_ using basic.stats function in *hierfstat* (71). We pooled each simulation into an empirical distribution of probable F_st_ values under the best fitting demographic model. We tested each observed site-wise F_st_ estimate against this empirical distribution to estimate a p-value. We corrected these p-values for multiple testing using a false discovery rate in R using the p.adjust function.

### Genes surrounding F_st_ outliers, GO term enrichment, and Variant Effects

We extracted the genes surrounding F_st_ outliers (p ≤ 0.05 after correction) in 10 Kb windows using the *D. pulicaria* RefSeq annotation (release 100, SC_F0-13Bv2) using bedtools (76). Next, we created Panther Generic Mappings for the genes with known annotations using each gene’s protein sequences following the method outlined in (41). Using the generic mappings, we tested for molecular function Gene Ontology (GO) term enrichment by testing against the *Daphnia pulex* gene list using the PantherDB webtool using a false discovery rate correction (41). In addition to testing for GO term enrichment, we also surveyed all genes for potential SNPs and small indels mutations potentially driving selection using Ensembl’s Variant Effect Predictor (VEP) (42). We built a custom database using the RefSeq annotations in GFF format following the developer’s protocol. We extracted all “High” (e.g., premature stops) and “Moderate” (e.g., missense SNPs) impact mutations predicted and cross-referenced these with the genes near F_st_ outliers.

### Chloride Tolerance

We selected a subset of 30 clones from the Lake (n = 10), 10-12 cm sediment layer (n =10), 18-20 cm (n = 3), and 22-24 cm (n = 7) subpopulations for estimating clone-specific tolerance to chloride. See the Supplemental Materials for details on the experimental set-up. We estimated LC_50_ for each clone separately by fitting a reduced-bias generalized linear model (GLM) to the survival curve using the R package *brglm* (77). Data were not normally distributed, nor did they meet the assumption of equal variances, so we performed a non-parametric ANOVA (Kruskal-Wallis) test on the LC_50_ estimates to test for differences in the mean LC_50_ value for each subpopulation.

## Acknowledgments

We would like to thank M. Edlund who facilitated the collection of the Tanners Lake sediment cores. T. Curb, Z. Arnold, A. Twumasi-Mensah, and A. Hemani, helped with the LC_50_ experiments. G. Wiley aided in the preparation of sequencing libraries. The University of Oklahoma Biological Station and the St. Croix Watershed Research Station facilitated field work. Funding for this study was provided by the University of Oklahoma Department of Biology Adams Summer Scholarship, Robberson Graduate College Grant, Hill Fund for Research in Biology, Graduate Student Senate Research Grant, AMNH Theodore Roosevelt Grant and the Biogeography of Behavior student seed grant (NSF DBI-2021880; PI L. Stein) awarded to MJW in support of graduate research. Any opinions, findings and conclusions or recommendations expressed in this material are those of the authors and do not necessarily reflect the views of the National Science Foundation, the American Museum of Natural History or the University of Oklahoma. This manuscript represents a portion of MJW’s doctoral dissertation at The University of Oklahoma. We thank three anonymous reviewers for constructive comments on earlier versions of the manuscript.

## Data Availability

Sequencing reads will be made available on the NCBI SRA upon acceptance. Data and appropriate metadata from this study will be archived in the University of Oklahoma’s ShareOK data repository (https://shareok.org). Code used in the analysis is available on GitHub at https://github.com/mwersebe/Tanner_lake_genomics.

## Notes

### Competing Interest Statement

The authors have declared no competing interest.

### Summary of Updates

Additional edits to the main text for clarity. Added new analysis and figures to the supplemental

