## Supplemental Figures for "Resurrection genomics provides molecular and phenotypic evidence of rapid adaptation to salinization in a keystone aquatic species"

Direct Submission Research Report

**Supplemental Methods:**

*Hatching Protocol:*

In brief, eggs (often paired) were excised from the ephippial casing using fine forceps and transferred into a well of a sterile 24-well cell culture plate filled with 2 mL of sterile COMBO media using a Pasteur pipette (1). Plates were covered in foil and incubated in a 4^o^ C refrigerator for 14 days and subsequently transferred to constant light at 20° C. Plates were checked daily for hatchlings for up to two weeks. Hatchlings as single individuals were carefully transferred to a 60 mL glass jar and fed a 50:50 mix of High Orthophosphate (HOP) COMBO grown *Scenedesmus* algae and diluted commercial *Nannochloropsis* concentrate (Reed Mariculture; 1 µL into 1 mL COMBO)*.*

*DNA Extraction, Library Preparation and Sequencing:*

Clones were expanded in 3.8 L jars so that approximately 100 adult females could be isolated. These 100 adults were subjected to a shortened antibiotic treatment described in (2). Genomic DNA was extracted from the resulting isoclonal pools (~50 adults per tube) using a MasterPure DNA extraction kit following the manufacturer’s directions (3). DNA was checked for quality by running a 2 µL aliquot on a 1% agarose electrophoresis gel and quantified using a Qubit fluorometer (ThermoFisher). Illumina-compatible sequencing libraries were constructed using the Ultra FS II DNA library preparation kit (New England Biolabs) and enriched using 4 cycles of PCR. Pooled and normalized libraries were sequenced across three separate runs of an Illumina NovaSeq instrument by the Oklahoma Medical Research Foundation. We sequenced the population to approximately 10X coverage on average (range 3- 20X).

*Chloride Tolerance:*

We reduced maternal effects among clones by raising experimental animals under standardized conditions. Ten to twenty gravid females per clone were isolated as single individuals into 60 mL jars filled with COMBO media from mass cultures (900 mL jars), for the establishment of experimental populations. These gravid females were allowed to release their clutches, the newborn animals (~25-35) were collected and isolated as single animals in fresh 60 mL jars and grown to adulthood and allowed to release their first clutches. Mothers were then transferred to fresh jars and allowed to release their second clutches. Animals from the second or third clutch for each clone that were born within 24 hours were isolated and pooled together into 900 mL jars and used in the experiments. We conducted 96-hour lethal concentration-50% assays (LC_50_) to estimate clone-specific tolerance to chloride. Experimental animals less than 24 hours old were haphazardly pipetted from the pooled (900-mL) jars and placed as single animals in 60 mL jars with COMBO media amended with either 0 mg (control), 250 mg, 500 mg, 1000 mg, or 1500 mg of Cl^-^ from NaCl. Each clone by concentration combination was replicated five times when animal numbers permitted (minimum was triplicate jars per treatment). Each jar across clones and treatments was assigned a random number and jars were placed in covered plastic shoe boxes in numerical order to reduce evaporative loss, as well as any bias due to positional effects. Experimental animals were maintained in a temperature-controlled room (20^o^ C) on a 12:12 light: dark cycle. We monitored animals daily (every 24 hours) for survival during the experiment. We recorded an animal as dead, when no movement of internal organs and/or appendages were observed after viewing under a dissecting microscope.

*mtDNA Analysis:*

To assess whether the TL population consisted of a single continuous population through time as the nuclear genome suggested, we analyzed the mtDNA of all clones by constructing a time-scaled phylogeny and inferred a Bayesian Skyline plot of the demographic history. Following the same bioinformatic approach outlined in the main text, we aligned reads to the *Daphnia pulex* mitochondrial reference genome (4). We called variants for each sample individually using BCFtools and applied SNPs variants to create a unique mitochondrial consensus sequence for each sample (5). We aligned each unique consensus sequence using MAFFT v7 (6). Next, we inferred a maximum likelihood phylogenetic tree in iqtree (7), and obtained branch support with 1000 ultra-fast bootstraps (8). We time-scaled out phylogeny using default parameters in TimeTree (9) and implemented a Bayesian Skyline analysis (10) to infer the demographic history from the mitochondrial sequences using BEAST2 (11). Tip dates were inferred from the sediment dating model proposed in Wersebe et al. (12), to the closest year (i.e., LC inferred to be from 2019, 22-24 cm from 1994, etc.).

*2021 Tanners Lake Diel Vertical Migration (DVM):*

On 29 June 2021 we returned to Tanners Lake to investigate the temporal and spatial heterogeneity of *Daphnia* occupancy of the water column. We quantitatively sampled the *Daphnia* population using a 30-L Schindler-Patalas trap at three stations covering the north-south axis of Tanners Lake. Beginning at 0600 hours (local time), we sampled the deep pelagic station (44.95076° N, -92.98135° W) trapping plankton at depths of 1-m, 6-m, 8-m and 12-m from on board an anchored canoe. We collected samples at a mid-depth pelagic station (44.95287° N, -92.98108° W) sampling the plankton density at 1-m, 6-m 9-m. Finally, we also collected samples from a shallow littoral station (44.95681° N, -92.98141° W) at 1-m and 2.5-m. In addition to the plankton samples, while at each station we also collected vertical profiles of temperature (°C), specific conductance (SPC; μS/cm) and dissolved oxygen (%) using a multiparamter sonde (YSI, Yellow Springs, OH) and the approximate Secchi depth (m).

We conducted two additional sampling bouts throughout the day, at 1200 hours and 2000 hours on 29 June 2021. During these times we only collected animals from the water column. Once on shore, we preserved animals according to the method outlined in Black and Dodson (14). Each sample was standardized to 50 mL of 70% EtOH. We estimated the number of collected *Daphnia* by subsampling each 50 mL sample 3 times with a 2 mL Hensen-Stempel Pipette and enumerating the animals over a grid. We identified animals to species using the key provided by Haney et al. (15). Using the average of the three Hensen-Stempel subsamples we calculated an estimate of the number of animals collected. Finally, this number was divided by 30-L (size of trap) to determine the density of animals (number per liter) present in the water column at the time of collection.

*2021 Tanners Lake DVM Study Results:*

We limit our DVM results here to just the focal species of our genetic study, *Daphnia pulicaria.* However, several other species were collected during our surveys. We detected *D. pulicaria* in every sample except from the shallow littoral sites (0600 2.5-m& 1200 1-m). In samples where we did detect *D. pulicaria,* densities ranged from a low of 0.185 individuals per liter (0600 12m deep) to a high of 27.225 animals per liter (0600 6 m deep). We consistently observed that the highest density estimates were at 6-m at the two pelagic sampling stations. Animals were observed at depth (12 m) throughout the day, ranging from 0.185 (0600) to 5.65 (1200) animals per liter. The density estimates of *D. pulicaria* are presented in Figure S6. Thermal and chemical stratification was much weaker in 2021 compared to our profiles taken in 2019 (see 12). The thermocline begins at approximately 3-m and ends at approximately 5.2-m of depth. SPC ranged from approximately 880 μS/cm at the surface (~190 mg/L Cl-) to a high at 13-m of 1108 μS/cm (~255 mg/L Cl-). Also, likely as a result of the weaker chemical stratification, DO was not completely depleted at depth and ranged from over 100% saturation at the surface to 5.1% saturation at 13-m (approximately 0.64 m/L O_2_). Temperature, SPC and DO profiles are available in Figure S7 A-C. Secchi depths at all stations ranged from ~2.5 m to 3.0 m and are available in Table S4.

**Supplemental Figures:**

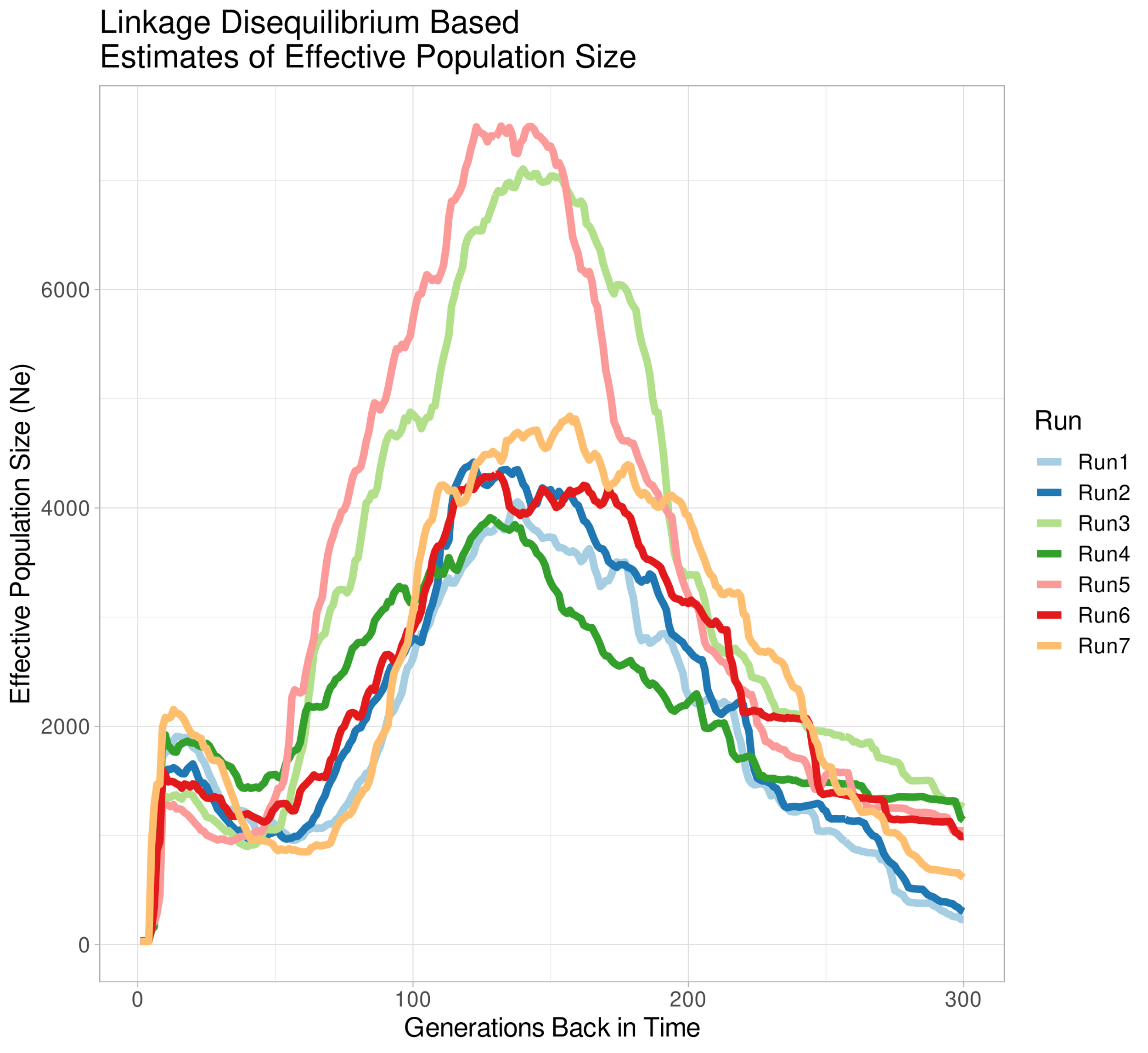

**Figure S1:** Effective population size (N_e_) trajectory inferred using Linkage Disequilibrium. The program GONE was run 7 random samples of 600,000 SNPs sampled from all the SNPs called in the Lake Clones (LC) subpopulation. Results were pruned to show the first 300 generations (method accurate up to ~200-300 generations back in time).

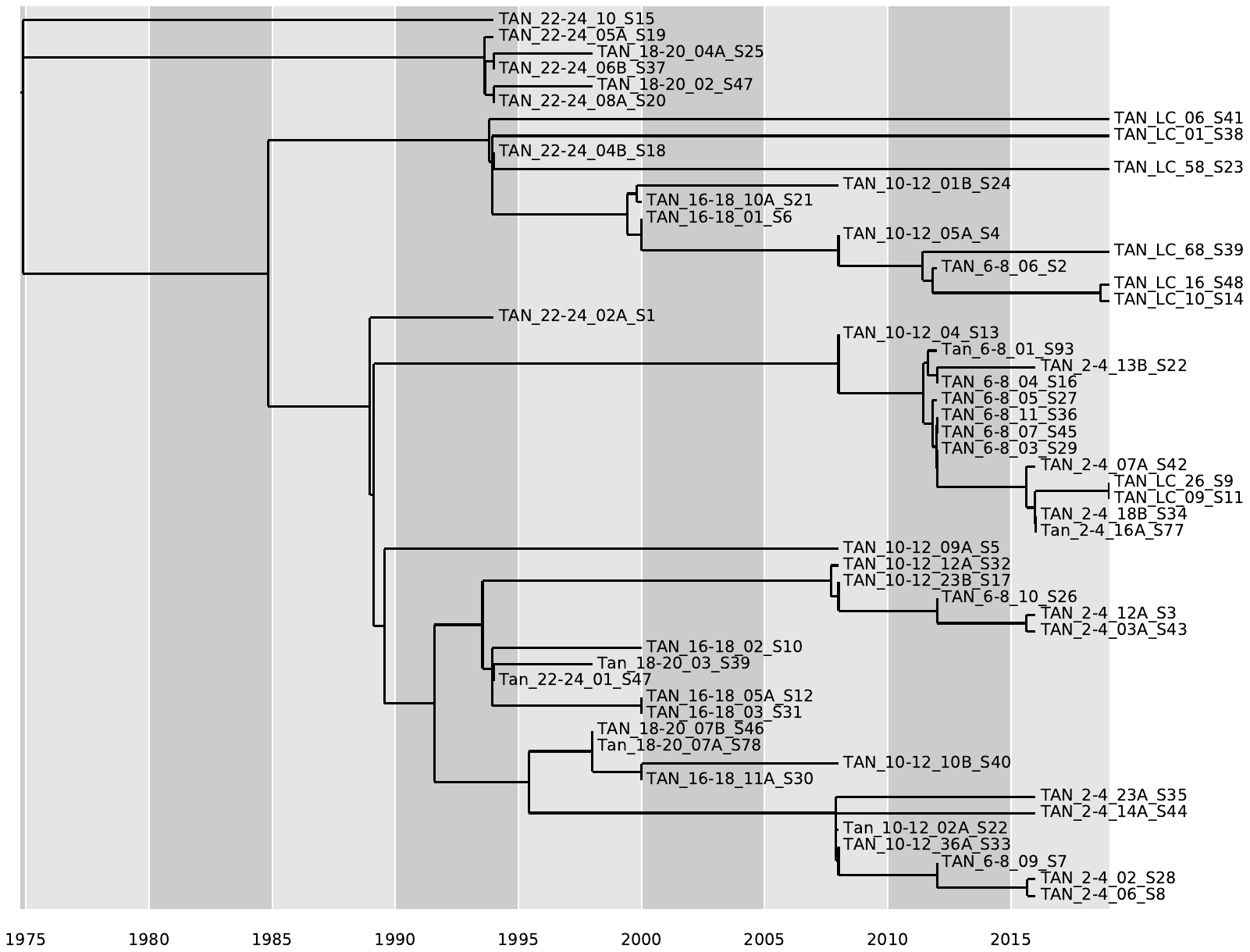

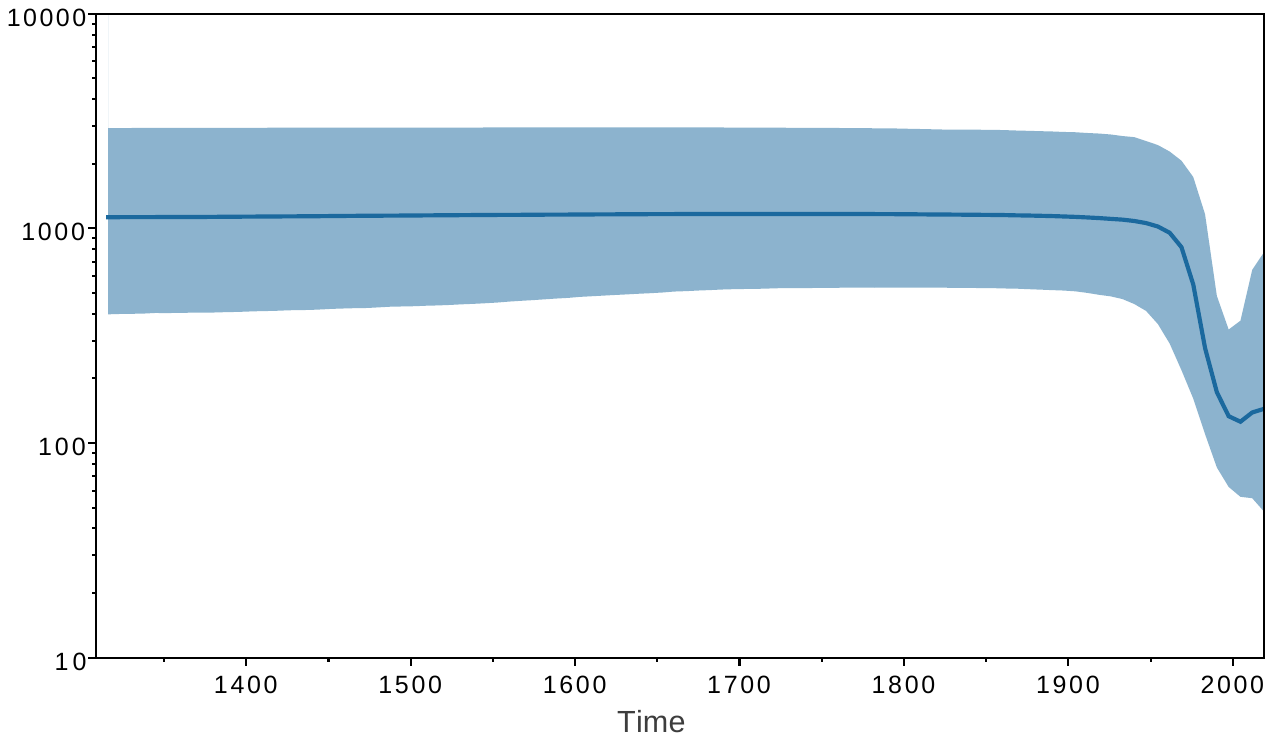

**Figure S2: (TOP)** Time-scaled maximum-likelihood phylogeny for full-length Tanners Lake (TL) mitochondrial genome sequences. Tip dates were assigned based on the age of the sediment layer of ephippial recovery inferred by the dating model for TL (see above) to nearest year. The tree was rooted with TimeTree to maximize the temporal signal using the default method. Nodes coalesced samples typically across time periods, indicating a constant population across time. **(BOTTOM)** Bayesian skyline plot inferred using BEAST2. The shape of the demographic history is similar to that inferred by GONE (see S1 above); however N_e_ sizes differed by an order of magnitude.

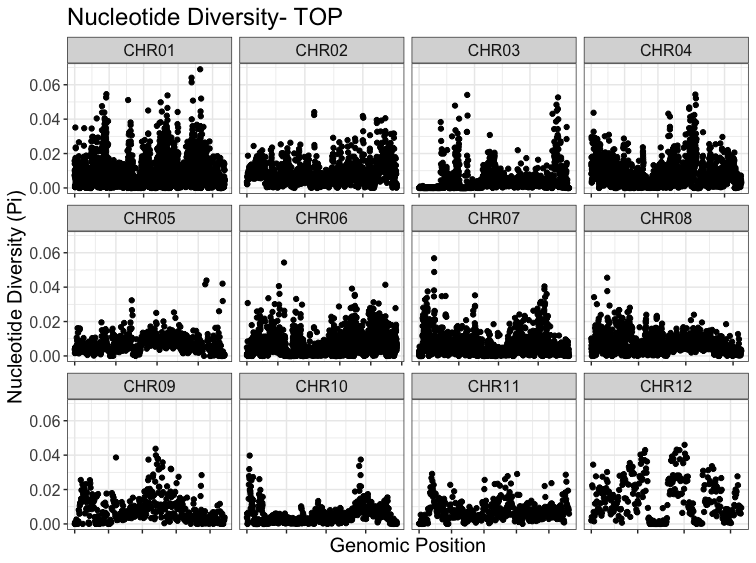

**Figure S3:** Windowed (10kb) nucleotide diversity genome-wide for the TOP population.

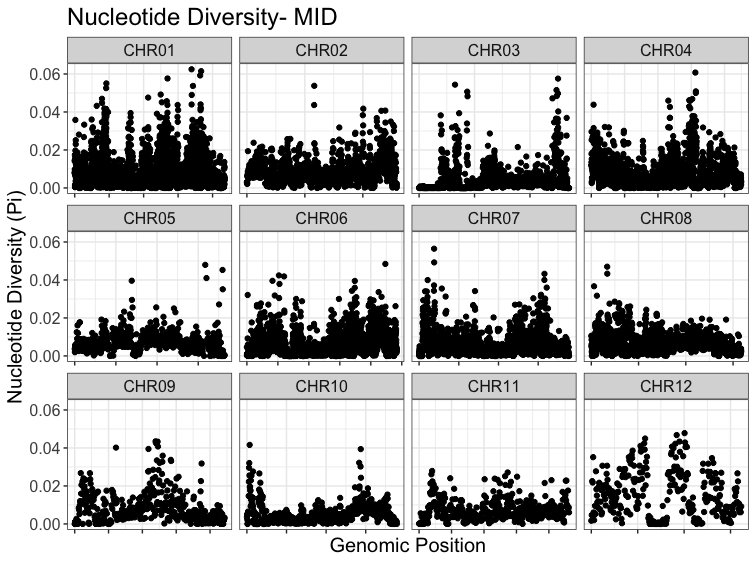

**Figure S4:** Windowed (10kb) nucleotide diversity genome-wide for the MID population.

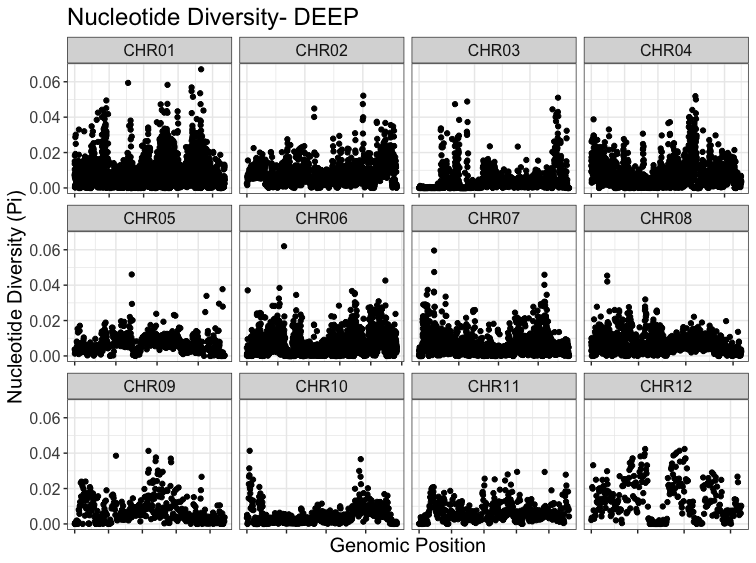

**Figure S5:** Windowed (10kb) nucleotide diversity genome-wide for the DEEP population.

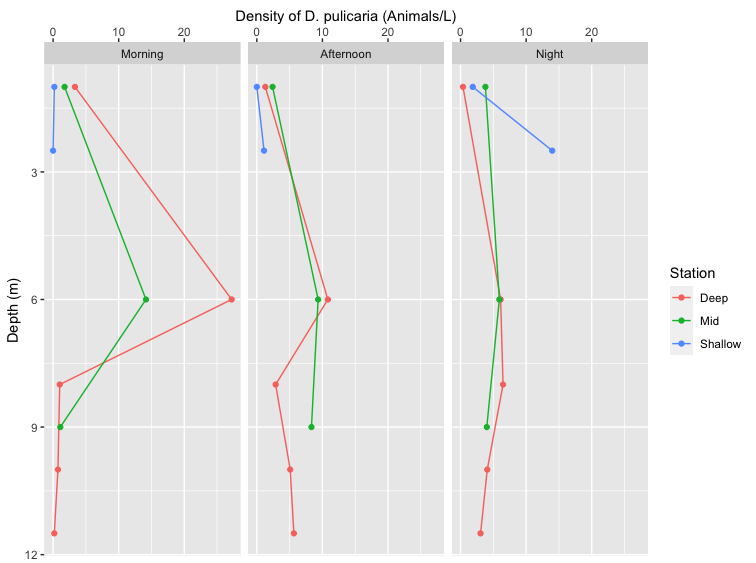

**Figure S6:** Spatial and temporal heterogeneity in *Daphnia pulicaria* densities. Colors represent the three sampling stations. Left Panel: *D. pulicaria* densities across depths at the three stations at 0600 hours. Middle Panel: *D. pulicaria* densities across depths at the three stations at 1200 hours. Right Panel: *D. pulicaria* densities across depths at the three stations at 2200 hours.

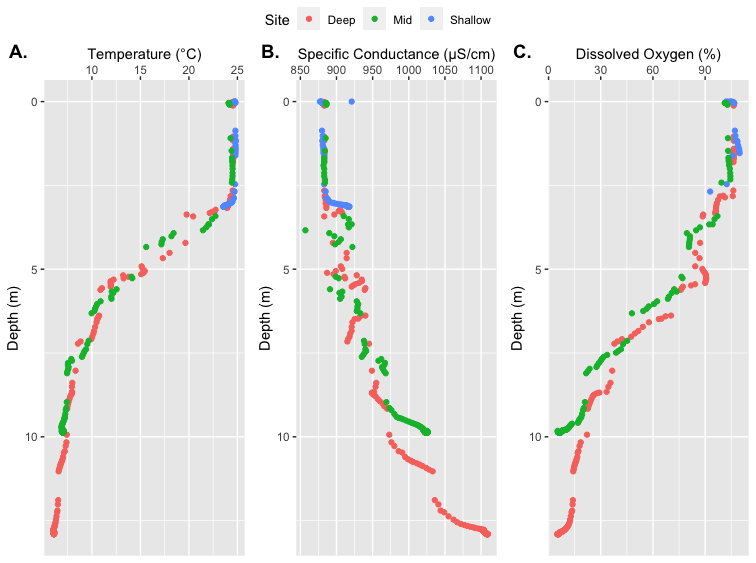

**Figure S7:** Abiotic profiles collected on 29 June 2021 collected with a YSI multiparameter sonde. Colors represent profiles taken at each sampling station. A) Temperature profile (°C). B) Specific conductance profile (μS/cm). C) Dissolved Oxygen profile (% saturation).

**Table S1:** 95% confidence intervals for the demographic model fit using FSC2.7.

|  | **Effective Population Size** | **Growth Rate** |
| --- | --- | --- |
| **Model** | 2931 | 2.6269 x 10^-5^ |
| **+ 95% Confidence** | 2939.162 | 2.6479 x 10^-6^ |
| **- 95% Confidence** | 2921.498 | -5.529 x 10^-6^ |
| **Mean** | 2930.33 | -1.44 x 10^-6^ |

**Table S2:** GO term enrichment analysis.

#Analysis Type:	PANTHER Overrepresentation Test (Released 20220202)

#Annotation Version and Release Date:	PANTHER version 17.0 Released 2022-02-22

#Analyzed List:	outlier_prots.panther.txt

#Reference List:	Daphnia pulex (all genes in database)

#Test Type:	FISHER

#Correction:	FDR

#PANTHER GO-Slim Molecular Function	Daphnia pulex - REFLIST (30047)	outlier_prots.panther.txt (201)	outlier_prots.panther.txt (expected)	outlier_prots.panther.txt (over/under)	outlier_prots.panther.txt (fold Enrichment)	outlier_prots.panther.txt (raw P-value)	outlier_prots.panther.txt (FDR)

protein kinase C binding (GO:0005080)	5	2	.03	+	59.80	9.03E-04	1.12E-02

voltage-gated potassium channel activity (GO:0005249)	14	3	.09	+	32.03	1.84E-04	3.47E-03

postsynaptic neurotransmitter receptor activity (GO:0098960)	11	2	.07	+	27.18	3.27E-03	3.27E-02

RNA polymerase II complex binding (GO:0000993)	12	2	.08	+	24.91	3.79E-03	3.72E-02

neuropeptide receptor activity (GO:0008188)	12	2	.08	+	24.91	3.79E-03	3.65E-02

3'-5'-exoribonuclease activity (GO:0000175)	13	2	.09	+	23.00	4.36E-03	4.04E-02

basal RNA polymerase II transcription machinery binding (GO:0001099)	13	2	.09	+	23.00	4.36E-03	3.97E-02

basal transcription machinery binding (GO:0001098)	13	2	.09	+	23.00	4.36E-03	3.90E-02

voltage-gated channel activity (GO:0022832)	33	5	.22	+	22.65	5.18E-06	5.28E-04

voltage-gated ion channel activity (GO:0005244)	33	5	.22	+	22.65	5.18E-06	4.40E-04

voltage-gated cation channel activity (GO:0022843)	27	4	.18	+	22.15	5.17E-05	1.47E-03

RNA polymerase core enzyme binding (GO:0043175)	14	2	.09	+	21.36	4.96E-03	4.29E-02

neuropeptide binding (GO:0042923)	15	2	.10	+	19.93	5.60E-03	4.76E-02

chloride channel activity (GO:0005254)	23	3	.15	+	19.50	6.71E-04	9.25E-03

potassium channel activity (GO:0005267)	35	4	.23	+	17.08	1.30E-04	3.01E-03

tetrapyrrole binding (GO:0046906)	29	3	.19	+	15.46	1.24E-03	1.44E-02

heme binding (GO:0020037)	29	3	.19	+	15.46	1.24E-03	1.41E-02

sulfotransferase activity (GO:0008146)	53	5	.35	+	14.10	4.24E-05	1.44E-03

gated channel activity (GO:0022836)	77	6	.52	+	11.65	1.97E-05	1.26E-03

potassium ion transmembrane transporter activity (GO:0015079)	53	4	.35	+	11.28	5.68E-04	8.04E-03

hydrolase activity, hydrolyzing O-glycosyl compounds (GO:0004553)	81	6	.54	+	11.07	2.58E-05	1.20E-03

hydrolase activity, acting on glycosyl bonds (GO:0016798)	88	6	.59	+	10.19	4.00E-05	1.46E-03

inorganic anion transmembrane transporter activity (GO:0015103)	46	3	.31	+	9.75	4.25E-03	4.02E-02

ion channel activity (GO:0005216)	145	8	.97	+	8.25	8.98E-06	6.54E-04

anion transmembrane transporter activity (GO:0008509)	76	4	.51	+	7.87	2.02E-03	2.10E-02

passive transmembrane transporter activity (GO:0022803)	163	8	1.09	+	7.34	2.01E-05	1.14E-03

channel activity (GO:0015267)	163	8	1.09	+	7.34	2.01E-05	1.03E-03

cation channel activity (GO:0005261)	110	5	.74	+	6.79	1.05E-03	1.27E-02

transmembrane signaling receptor activity (GO:0004888)	168	7	1.12	+	6.23	1.78E-04	3.48E-03

peptide binding (GO:0042277)	98	4	.66	+	6.10	4.84E-03	4.26E-02

inorganic molecular entity transmembrane transporter activity (GO:0015318)	210	8	1.40	+	5.69	1.12E-04	2.72E-03

RNA polymerase II cis-regulatory region sequence-specific DNA binding (GO:0000978)	215	8	1.44	+	5.56	1.31E-04	2.90E-03

ion transmembrane transporter activity (GO:0015075)	242	9	1.62	+	5.56	4.99E-05	1.50E-03

cis-regulatory region sequence-specific DNA binding (GO:0000987)	216	8	1.44	+	5.54	1.35E-04	2.86E-03

signaling receptor activity (GO:0038023)	279	10	1.87	+	5.36	2.58E-05	1.10E-03

molecular transducer activity (GO:0060089)	280	10	1.87	+	5.34	2.66E-05	1.04E-03

DNA-binding transcription factor activity (GO:0003700)	324	9	2.17	+	4.15	4.14E-04	6.40E-03

RNA polymerase II transcription regulatory region sequence-specific DNA binding (GO:0000977)	330	9	2.21	+	4.08	4.71E-04	7.07E-03

DNA-binding transcription factor activity, RNA polymerase II-specific (GO:0000981)	304	8	2.03	+	3.93	1.21E-03	1.43E-02

transcription cis-regulatory region binding (GO:0000976)	350	9	2.34	+	3.84	7.09E-04	9.51E-03

transcription regulatory region nucleic acid binding (GO:0001067)	350	9	2.34	+	3.84	7.09E-04	9.27E-03

phosphotransferase activity, alcohol group as acceptor (GO:0016773)	313	8	2.09	+	3.82	1.45E-03	1.57E-02

double-stranded DNA binding (GO:0003690)	393	10	2.63	+	3.80	3.92E-04	6.24E-03

transcription regulator activity (GO:0140110)	438	11	2.93	+	3.75	2.25E-04	3.83E-03

sequence-specific double-stranded DNA binding (GO:1990837)	361	9	2.41	+	3.73	8.77E-04	1.12E-02

transmembrane transporter activity (GO:0022857)	407	10	2.72	+	3.67	5.11E-04	7.45E-03

DNA binding (GO:0003677)	579	14	3.87	+	3.61	4.72E-05	1.51E-03

kinase activity (GO:0016301)	346	8	2.31	+	3.46	2.65E-03	2.71E-02

sequence-specific DNA binding (GO:0043565)	391	9	2.62	+	3.44	1.50E-03	1.60E-02

| **Chromosome** | **Position** | **Fst** | **Type** | **Gene** |
| --- | --- | --- | --- | --- |
| 04 | 13394600 | 0.3983 | Intronic | *pank4; tda6* |
| 02 | 6974770 | 0.3645 | Intergenetic | NA |
| 02 | 3614856 | 0.3635 | Intronic | *col1a1; tcaim* |
| 03 | 8229625 | 0.3609 | Intronic | *rap1gap* |
| 03 | 7485508 | 0.3480 | Intronic | *daam1* |
| 05 | 4372154 | 0.3284 | Intronic | *ptn* |
| 06 | 21993985 | 0.3221 | Intronic | *rotund* |
| 04 | 2181789 | 0.3220 | Intronic | *still life* |
| 04 | 19599247 | 0.3211 | Intronic | *fstl5* |
| 05 | 4380601 | 0.3206 | Intronic | *ptn* |
| 02 | 9684091 | 0.3175 | Intronic | Uncharacterized:  LOC124326411  LOC124326443  LOC124326498 |

**Table S3:** Extreme outlier SNPs and nearest genes (within 5Kb).

**Table S4:** Secchi Depths at sampling stations on Tanners Lake

| **Time** | **0600** | **1200** | **2000** |
| --- | --- | --- | --- |
| **Deep Pelagic** | 3.0 m | 2.9 m | 2.9 m |
| **Mid Pelagic** | 3.0 m | 2.85 m | 2.8 m |
| **Shallow Littoral** | 3.1 m | 2.9 m | 2.9 m |
